# Ultra-performance liquid chromatography-mass spectrometry analysis of post-mortem brain tissue reveals specific amino acid profile dysregulation in Parkinson’s Disease and Alzheimer’s Disease patients

**DOI:** 10.1101/2025.07.22.666071

**Authors:** Jacopo Gervasoni, Anna Di Maio, Marcello Serra, Michela Cicchinelli, Lavinia Santucci, Gabriele Ciasca, Tommaso Nuzzo, Qin Li, Marie-Laure Thiolat, Micaela Morelli, Andrea Urbani, Francesco Errico, Erwan Bezard, Alessandro Usiello

## Abstract

**Background:** Combined metabolomic and HPLC-based analyses have identified significant metabolic alterations in serum and plasma amino acid levels of Parkinson’s disease (PD) patients, underscoring their potential as biomarkers. However, it remains unclear whether these biochemical changes also manifest within the central nervous system or are confined to peripheral metabolism, reflecting systemic metabolic disturbances.

**Methods:** To address this issue, here we measured the levels of 44 different amino acids in *post-mortem* brain samples from MPTP-intoxicated, L-DOPA-treated monkeys and PD patients at different Braak Lewy body (LB) stages, compared to their respective controls, through targeted Ultra-performance liquid chromatography–mass spectrometry (UPLC-MS).

**Results:** In MPTP-intoxicated monkeys, UPLC-MS revealed significant elevations in GABA, citrulline, threonine, isoleucine, phenylalanine, valine, glycine, and serine in the putamen, whereas we failed to detect alterations in the superior frontal gyrus (SFG). In PD patients, caudate-putamen (CPu) analysis demonstrated consistent serine upregulation across Braak LB stages 3–4 and 6, with stage 6 specifically showing additional proline increases and phosphoethanolamine decreases. Notably, serine was the sole amino acid significantly altered in both the putamen of MPTP-intoxicated monkeys and the CPu of PD patients. No significant amino acid alterations were observed in the SFG of PD patients, mirroring the findings in monkeys. In contrast, Alzheimer’s disease (AD) patient SFG samples showed significant increases in tryptophan, phenylalanine, threonine, tyrosine, and methionine relative to controls.

**Conclusions:** These findings demonstrate that cerebral amino acid alterations in PD are region-specific and primarily localized to brain areas receiving nigrostriatal dopaminergic innervation. Moreover, the cortical amino acid profile in AD differs substantially from that in PD, suggesting disease-specific metabolic signatures in distinct neurodegenerative conditions.

## 1. Background

Parkinson’s disease (PD) is the second most prevalent neurodegenerative disorder after Alzheimer’s disease (AD), affecting approximately 1-3% of individuals aged 60 years and older [1]. Notably, the global prevalence of PD is rising significantly, with a particularly pronounced increase observed over the past two decades [1]. Neuropathologically, PD is characterized by the loss of dopaminergic neurons in the substantia nigra pars compacta and the presence of Lewy bodies and neurites— abnormal aggregates of insoluble proteins primarily composed of α-synuclein (α-syn) [2].

PD is increasingly recognized as a multifactorial, systemic, and heterogeneous disorder, characterized by marked variability in motor symptoms (tremor, rigidity, bradykinesia, postural instability), non-motor symptoms (hyposmia, autonomic dysfunction, sleep disturbances, cognitive impairment), and treatment response [3]. This heterogeneity poses significant challenges to clinical diagnosis— primarily dependent on clinical and neuroimaging evaluation in the 85–90% of idiopathic cases — and for treatment selection [4].

To disentangle this complex scenario, a series of putative biomarkers, encompassing clinical indicators, neuroimaging, genomics, and biomolecules, has been proposed to enhance diagnostic accuracy and monitor disease progression [5]. Notwithstanding the valuable insights these biomarkers have provided regarding aspects of patients’ pathological conditions, none have yet demonstrated the specificity required to definitively diagnose PD, track disease progression, or predict treatment response [5,6].

Over the last decade, metabolomic investigation—through the comprehensive analysis of small molecules or metabolites within a biological system— has emerged as a powerful approach for elucidating the complex biochemical alterations associated with disease states. Studies conducted across different biological matrices, including serum, plasma, urine, cerebrospinal fluid (CSF), and saliva, in both animal models of PD and Parkinsonian patients, have identified numerous systemic metabolic alterations that provide valuable insights into the disease’s pathophysiology [7,8]. These alterations encompass dysregulation in lipid homeostasis [9–13], mitochondrial energy metabolism [10,14–16], ammonium recycling and purine biosynthesis pathways [17–19].

Additionally, metabolomic and High-Performance Liquid Chromatography (HPLC) investigations have revealed significant alterations in the homeostasis of neuroactive D– and L-amino acids in blood serum/plasma and CSF samples from patients with PD compared to controls [14,20–26]. Altered amino acid profiles in PD patients are thought to reflect disruptions in protein biosynthesis, neurotransmitter homeostasis, and key cellular signalling pathways, including urea cycle function, antioxidant (e.g., glutathione) synthesis, and energy metabolism [23]. Importantly, from a multi-organ perspective, these amino acid alterations may provide critical insights into disease progression by revealing metabolic dysregulation in peripheral organs—such as the kidneys and liver—in patients with PD [27,28].

While these biochemical alterations underscore the potential for developing novel “fingerprints” for PD, they also reveal the inherent complexity of PD pathophysiology. In this regard, conflicting results and inconsistencies across studies render it challenging to draw definitive conclusions regarding the central molecular events driving these changes [29].

A key challenge that complicates mechanistic interpretation of available data is that most of the above cited metabolomic studies have been conducted using patients’ peripheral biofluids, particularly plasma and serum, whereas investigations of brain tissue, particularly in the caudate-putamen (CPu), the basal ganglia nucleus most affected by dopaminergic nigrostriatal denervation, remain considerably more limited.

For instance, it’s unclear whether the distinct PD signature previously identified in patients’ blood also occurs centrally. Factors such as blood-brain barrier permeability, dietary influences, pharmacological treatments, systemic metabolism, and comorbid conditions can profoundly reshape peripheral metabolomic profiles, thereby complicating accurate extrapolation to the brain’s metabolic state.

To clarify this outstanding issue, the present study aims to employ a targeted metabolic approach using UPLC-MS—the gold standard in clinical chemical diagnostics—to quantify and compare the levels of 44 distinct amino acids *in post-mortem* samples obtained from: (1) the rostral putamen and superior frontal gyrus (SFG) of 1-methyl-4-phenyl-1,2,3,6-tetrahydropyridine (MPTP)-intoxicated, L-DOPA-treated macaques and their respective controls; (2) the CPu and SFG of PD patients at different Braak Lewy body (LB) stages and matched controls; and (3) the SFG of patients with Alzheimer’s disease (AD) and matched controls. Our metabolomic analysis revealed significant amino acid profile alterations in *post-mortem* striatal—but not cortical—samples from both Parkinsonian monkeys and PD patients compared to controls. These amino acid changes only partially overlap between species and, in PD patients, were restricted to serine, proline, and phosphoethanolamine. Lastly, we found multiple amino acid alterations in the SFG of AD patients compared to controls, suggesting a prominent role of underlying neurodegenerative pathology in cortical amino acid profile

## 2. Methods

### 2.1. Human post-mortem tissue collection

*Post-mortem* tissue samples from the caudate-putamen (CPu) of Parkinson’s disease (PD) patients and superior frontal gyrus (SFG) of PD and Alzheimer’s disease (AD) patients were obtained from The Netherlands Brain Bank (Netherlands Institute for Neuroscience, Amsterdam; open access: www.brainbank.nl) and derived from three distinct cohorts of patients and corresponding non-demented control subjects. Specifically, CPu tissues derived from a cohort of 26 PD patients comprising 13 individuals classified as Braak Lewy body (LB) stage 3–4 and 13 individuals as Braak LB stage 6. Due to the limited availability of *post-mortem* CPu samples from controls, PD patients were compared with 6 non-demented controls matched for age, sex, and *post-mortem* interval (PMI) to minimize potential confounding effects in the statistical analysis. *Post-mortem* SFG samples derived from a cohort of subjects consisting of patients with PD, AD, and controls. The number of PD patients (Braak LB stage ≥4; n = 10) was comparable to that from non-demented controls (n = 10) as well as the number of AD patients (neuropathological staging of Braak ≥5; n=10) and both PD and AD groups were matched with the control group for age, sex, and PMI. Non-demented controls were selected from adults with no cognitive decline and with a Braak LB score ≤3 in accordance with the Braak and Braak criteria [30]. Demographic characteristics of PD patients and non-demented controls are described in Table 1 (CPu). Demographic characteristics of the other cohorts, including patients with PD, AD and non-demented controls, are described in Table 2 (SFG). Control subjects had no known clinical history of neurological or psychiatric disorders, as confirmed by neuropathological evaluation of their samples. The clinical diagnosis of PD was established according to the UK Brain Bank criteria [31], subsequently confirmed by neuropathological findings [32] and AD was diagnosed based on National Institute of Neurological and Communicative Disorders and Stroke and the Alzheimer’s Disease and Related Disorders Association (NINCDS-ADRDA) criteria [33]. Frozen CPu and SFG tissues were pulverized in liquid nitrogen and stored at –80 °C for subsequent processing.

**Table 1.**
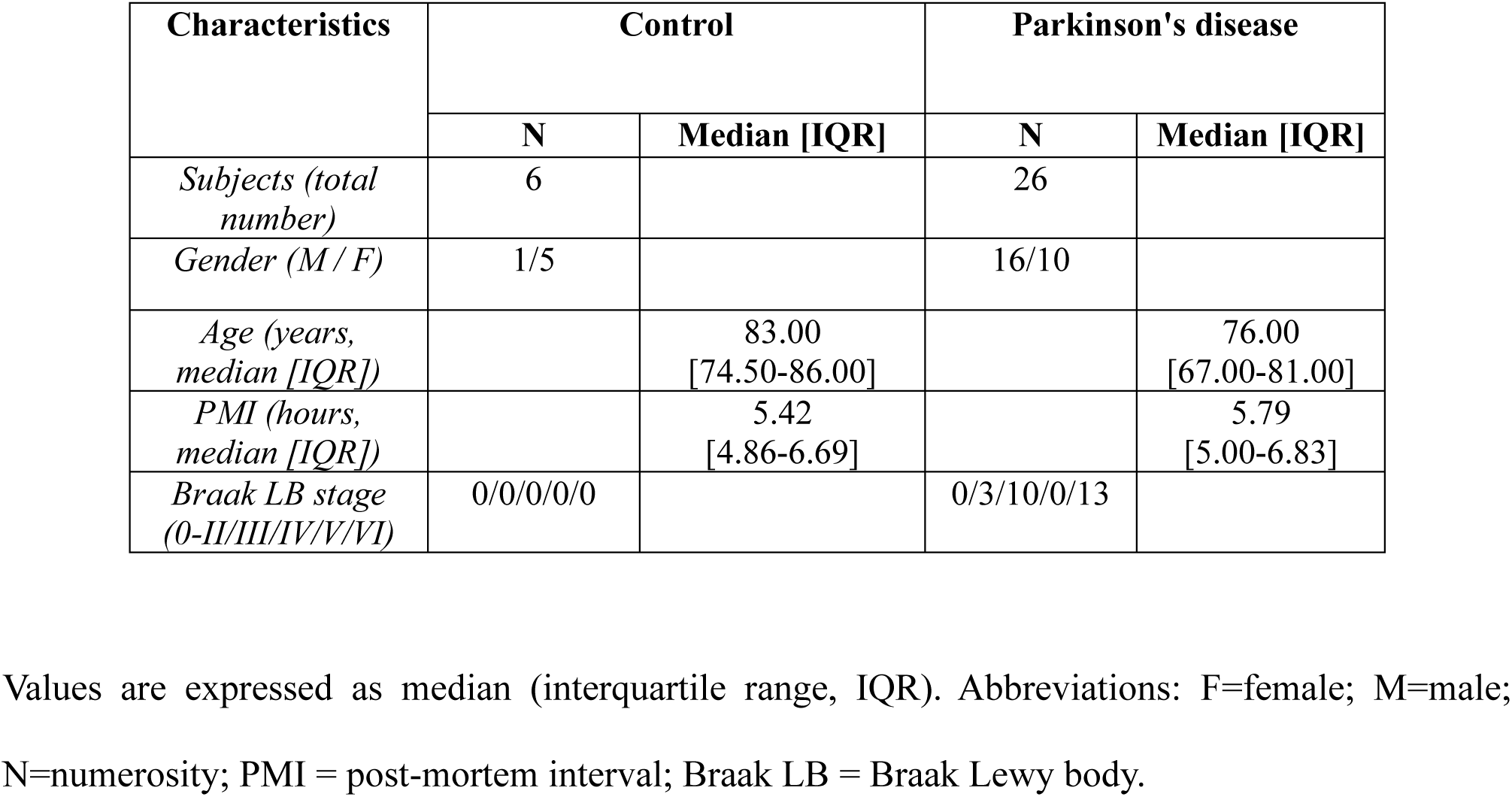
Demographic and clinical characteristics of Parkinson’s disease patients and non-demented control subjects from *post-mortem* caudate-putamen samples.

| Characteristics | Control |  | Parkinson's disease |  |
| --- | --- | --- | --- | --- |
|  | N | Median [IQR] | N | Median [IQR] |
| *Subjects (total number)* | 6 |  | 26 |  |
| *Gender (M / F)* | 1/5 |  | 16/10 |  |
| *Age (years, median [IQR])* |  | 83.00<br>[74.50-86.00] |  | 76.00<br>[67.00-81.00] |
| *PMI (hours, median [IQR])* |  | 5.42<br>[4.86-6.69] |  | 5.79<br>[5.00-6.83] |
| *Braak LB stage (0-II/III/IV/V/VI)* | 0/0/0/0/0 |  | 0/3/10/0/13 |  |

Values are expressed as median (interquartile range, IQR). Abbreviations: F=female; M=male; N=numerosity; PMI = post-mortem interval; Braak LB = Braak Lewy body.

**Table 2.** Demographic and clinical characteristics of Alzheimer’s disease, Parkinson’s disease patients and non-demented control subjects from *post-mortem* superior frontal gyrus samples.

| Characteristics | Control |  | Alzheimer's disease |  | Parkinson's disease |  |
| --- | --- | --- | --- | --- | --- | --- |
|  | N | Median [IQR] | N | Median [IQR] | N | Median [IQR] |
| *Subjects (total number)* | 10 |  | 10 |  | 10 |  |
| *Gender (M / F)* | 10/0 |  | 10/0 |  | 10/0 |  |
| *Age (years, median [IQR])* |  | 85.00<br>[79.00-88.25] |  | 85.50<br>[82.00-88.00] |  | 85.00<br>[81.75-86.00] |
| *PMI (hours, median [IQR])* |  | 5.87<br>[5.62-6.58] |  | 5.37<br>[5.04-6.42] |  | 5.79<br>[4.42-6.45] |
| *Braak stage (0-I/II-IV/V-VI)* | 3/7/0 |  | 0/0/10 |  |  |  |
| *Braak LB stage*<br>*(0-II/III/IV/V/VI)* | 4/0/0/0/0 |  |  |  | 0/0/2/2/6 |  |

Values are expressed as median (interquartile range, IQR). Abbreviations: F=female; M=male; N= numerosity; PMI = post-mortem interval; Braak LB = Braak Lewy body.

### 2.2 Non-human primates

Tissues used in the present study were obtained from an experimental brain bank, with detailed protocols described elsewhere [34–38]. Briefly, 1-methyl-4-phenyl-1,2,3,6-tetrahydropyridine (MPTP)-intoxicated non-human primate PD model macaques (n = 5) received daily intravenous injections of MPTP hydrochloride (0.2 mg/kg) until stable parkinsonian motor symptoms developed. Subsequently, animals were administered individually titrated doses of L-DOPA (Madopar; L-DOPA/carbidopa, 4:1 ratio; range: 9–17 mg/kg), twice daily, to achieve optimal reversal of motor deficits. This treatment was maintained for 4–5 months until dyskinesia stabilized. Thereafter, L-DOPA was administered biweekly to sustain a consistent dyskinetic state prior to acute pharmacological testing. At the end of the study, animals were euthanized via intravenous overdose of sodium pentobarbital (150 mg/kg) one hour after the final administration of L-DOPA, and brains were rapidly extracted. A control group of five macaques, not treated with MPTP hydrochloride or L-DOPA, was included as control. Each brain was bisected along the midline, and both hemispheres were immediately frozen in isopentane at −45 °C, then stored at −80 °C. Coronal sections (300 μm thick) were cut using a cryostat, and tissue punches were obtained from the rostral putamen and the SFG. As previously reported [37], this experimental procedure induces a significant reduction (>75%) in striatal tyrosine hydroxylase protein expression and dopamine levels.

### 2.3 Ultra-performance liquid chromatography-mass spectrometry

Brain tissue samples from humans and monkeys were homogenized in 1:20 (w/v) 0.2 M trichloroacetic acid (TCA), sonicated (3 cycles, 10 s each) and centrifuged at 13,000 × g for 20 min. The TCA supernatants were used for subsequent analyses.

The concentrations of a panel of 44 amino acids and derivatives were measured in brain tissue extracts from Parkinsonian monkeys, patients with PD and AD, and their respective matched controls, using UPLC-MS. The panel includes: 1-Methylhistidine, 3-Methylhistidine, 4-Hydroxyproline, α-Aminobutyric acid, β-Alanine, β-Aminobutyric acid, γ-Aminobutyric acid, Alanine, Allo-Isoleucine, Aminoadipic acid, Anserine, Arginine, Asparagine, Aspartic acid, Carnosine, Citrulline, Cystathionine, Cystine, Ethanolamine, Glutamic acid, Glutamine, Glycine, Glycil proline, Histidine, Homocitrulline, Homocystine, Hydroxylysine, Isoleucine, Kynurenine, Leucine, Lysine, Methionine, Ornithine, Phenylalanine, Phosphoethanolamine, Proline, Sarcosine, Serine, Sulfocysteine, Taurine, Threonine, Tryptophan, Tyrosine, Valine.

Sample preparation was performed adding 50 µL of the sample to 100 µL 10% (w/v) sulfosalicylic acid containing an internal standard mix (50 µM) (Cambridge Isotope Laboratories, Inc., Tewksbury, MA, USA). The mixture was vortexed and then centrifuged at 10,000 rpm for 15 min. In a 1.5 mL vial, 70 µL of borate buffer and 20 µL of AccQ Tag reagents (Waters Corporation, Milford, MA, USA) were added to 10 µL of the obtained supernatant and heated at 55 °C for 10 min in a water bath. Next, samples were loaded onto a CORTECS UPLC C18 column 1.6 µm, 2.1 mm x 150 mm (Waters Corporation) for chromatographic separation (ACQUITY H-Class, Waters Corporation). Elution was accomplished at 0,5 mL/min flow-rate with a linear gradient (9 min) from 99:1 to 1:99 water 0.1% formic acid/acetonitrile 0.1% formic acid. Injection volume was 2 µL and column oven temperature was set at 55°C. Analytes were detected on an ACQUITY QDa single quadrupole mass spectrometer equipped with an electrospray source operating in positive ion mode (Waters Corporation).

The analytical process was monitored using amino acids controls (level 1 and level 2) manufactured by the MCA laboratory of the Queen Beatrix Hospital (The Netherlands). Amino acids concentrations were determined by comparison with values obtained from a standard curve for each amino acid (2,5-10-50-125-250-500 μmol/L only for Cysteine, 5-20-100-250-500-1000 μmol/L for all amino acids). For data analysis (calibration curves and amino acid quantitation), the instrument software TargetLynx was used.

### 2.4 Statistical analysis

UPLC/MS data were processed to construct four datasets. The first dataset (Monkeys’ putamen) was obtained from the *post-mortem* putamen of MPTP-intoxicated, L-DOPA-treated monkeys (n=5) and untreated controls (n=5). The second dataset (Monkeys’ SFG) was built from the *post-mortem* SFG of MPTP-intoxicated, L-DOPA-treated monkeys (n=5) and untreated controls (n=5). The third dataset (human CPu) was obtained from the *post-mortem* CPu of PD patients (n=26) and non-demented controls (n=6). The CPu samples of PD patients were further divided based on the Braak LB stages (3–4 and 6). The fourth dataset (Human SFG) was built from the *post-mortem* SFG of PD patients (n=10), AD patients (n=10), and non-demented controls (n=10). The SFG samples of PD patients were further divided based on the Braak LB stages (4–5 and 6). The following conditions were applied for variables (amino acids) inclusion in every dataset: amino acids that were not detected or had more than 30% of missing values (m.v.) were excluded. The remaining m.v. were then imputed using the Probabilistic Principal Component Analysis (PPCA) method. The final datasets were set up as follows:

- Monkeys’ putamen dataset with p=24 variables and n=10 total samples (0.3% imputed m.v.),
- Monkeys’ SFG dataset with p=22 variables and n=10 total samples (1.2% imputed m.v.),
- Human CPu dataset with p=27 variables and n=33 total samples (1% imputed m.v.),
- Human SFG dataset with p=25 variables and n=30 total samples (1.5% imputed m.v.).

All the quantified amino acids were first normalized for protein concentration (nmol/mg). Then, datasets were normalized according to the specific data distribution. The Monkeys’ putamen and Monkeys’ SFG datasets were log-transformed and auto scaled, while the Human CPu and Human SFG datasets were normalized by median and Pareto scaling. Differential amino acid levels were obtained using a Fold Change (FC) threshold set at 1.2. Statistical significance was assessed using t-test with significance defined as p-value < 0.05, or Analyses of Variance (ANOVA) with a p-value < 0.05, and Tukey’s HSD post hoc with False Discovery Rate (FDR) < 0.05, reporting the degrees of freedom (DF) for every analysis. Volcano plots were created using FC = 1.2 and p-value < 0.05 as thresholds. All statistical analyses, volcano plots, and violin plots were generated using MetaboAnalyst 6.0 (http://www.metaboanalyst.ca/). Clustergrams were obtained using R (Version 4.4.2, R Core Team, R Foundation for Statistical Computing, Vienna, Austria) with the pheatmap, dplyr, and ggplot2 packages. The Venn diagram was created using PowerPoint 2013.

## 3. Results

### Amino Acid Levels Are Site-Specifically Altered in MPTP-intoxicated, L-DOPA-treated Monkeys

First, utilizing the MPTP macaque model, considered a gold standard preclinical model of PD [39], we investigated the impact of severe nigrostriatal dopamine degeneration and repeated L-DOPA treatment on the putaminal and cortical concentrations of a panel of 44 amino acids. Specifically, a univariate statistical analysis was performed across the Monkeys’ putamen and Monkeys’ SFG datasets, with the aim to identify differentially expressed amino acids in MPTP-intoxicated, L-DOPA-treated monkeys compared to non-treated controls in the two brain regions. A two-sample t-test was employed to assess statistical significance, while fold change (FC) analysis was used to quantify the degree and direction of alterations in amino acid levels. In the putamen, the analysis revealed a distinct pattern of metabolic dysregulation, with eight amino acids found to be significantly upregulated in MPTP-intoxicated, L-DOPA-treated monkeys compared to the controls: serine (FC 2.08; p-value 8.19×10^−4^), glycine (FC 1.64; p-value 0.014), threonine (FC 1.86; p-value 0.027), gamma-amino butyric acid (GABA) (FC 1.70; p-value 0.015), phenylalanine (FC 1.52; p-value 0.016), valine (FC 1.60; p-value 0.037), isoleucine (FC 1.57; p-value 0.037), and citrulline (FC 1.63; p-value 0.019) (Fig. 1).

**Figure 1.**
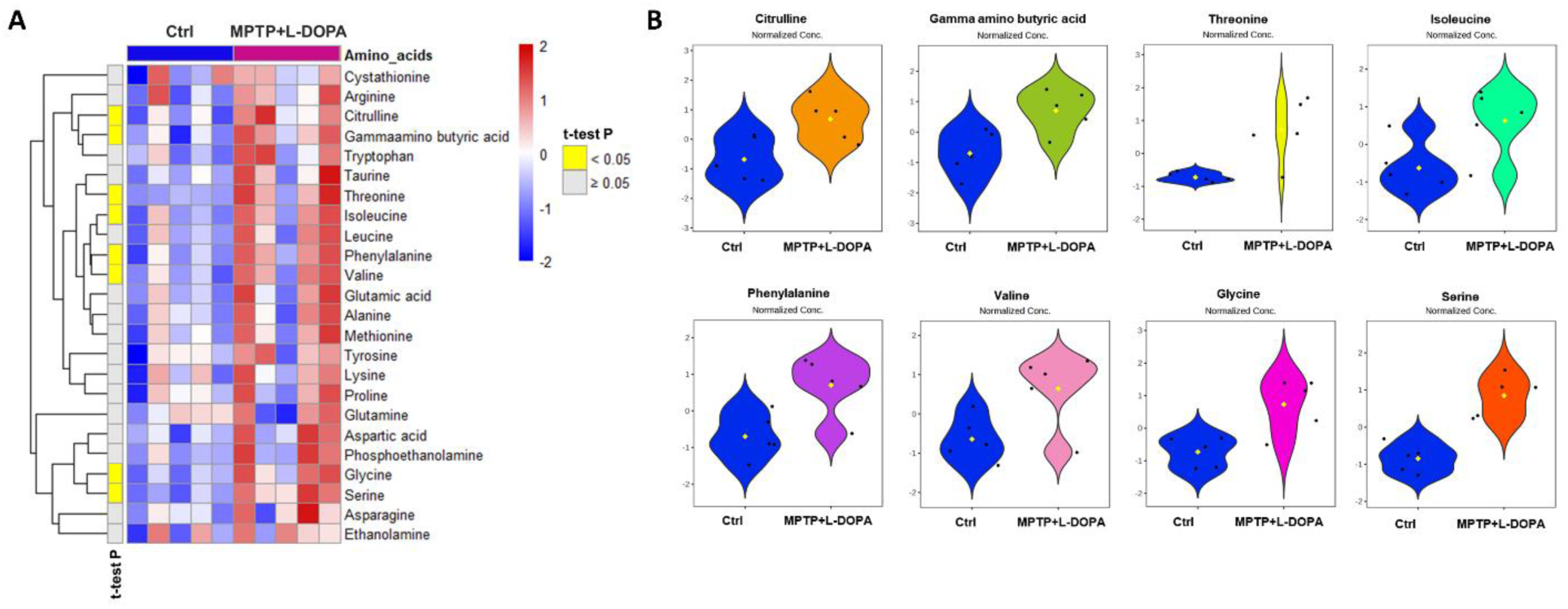
MPTP-intoxicated, L-DOPA-treated monkeys showed upregulated putaminal levels of serine, glycine, threonine, GABA, phenylalanine, valine, isoleucine, and citrulline. A) The clustergram displays amino acids within the Monkeys’ putamen dataset. Rows represent amino acids (variables), while columns the samples, divided into control Ctrl (n=5) and MPTP+L-DOPA (n=5). The elements of the clustergram illustrate amino acid concentrations in samples, standardized using a z-score, visualized using a color scale ranging from blue (lower) to red (higher). The first column indicates t-test statistical significance, showing significant amino acids in yellow and not significant ones in gray. B) Violin plots depict the concentrations of amino acids that differ significantly between Ctrl and MPTP+L-DOPA monkeys. Abbreviations: Ctrl, control; MPTP, 1-methyl-4-phenyl-1,2,3,6-tetrahydropyridine.

In contrast, no significant alterations in amino acid concentrations were detected in the *post-mortem* SFG of MPTP-intoxicated, L-DOPA-treated monkeys compared to the controls (Fig. 2).

**Figure 2.**
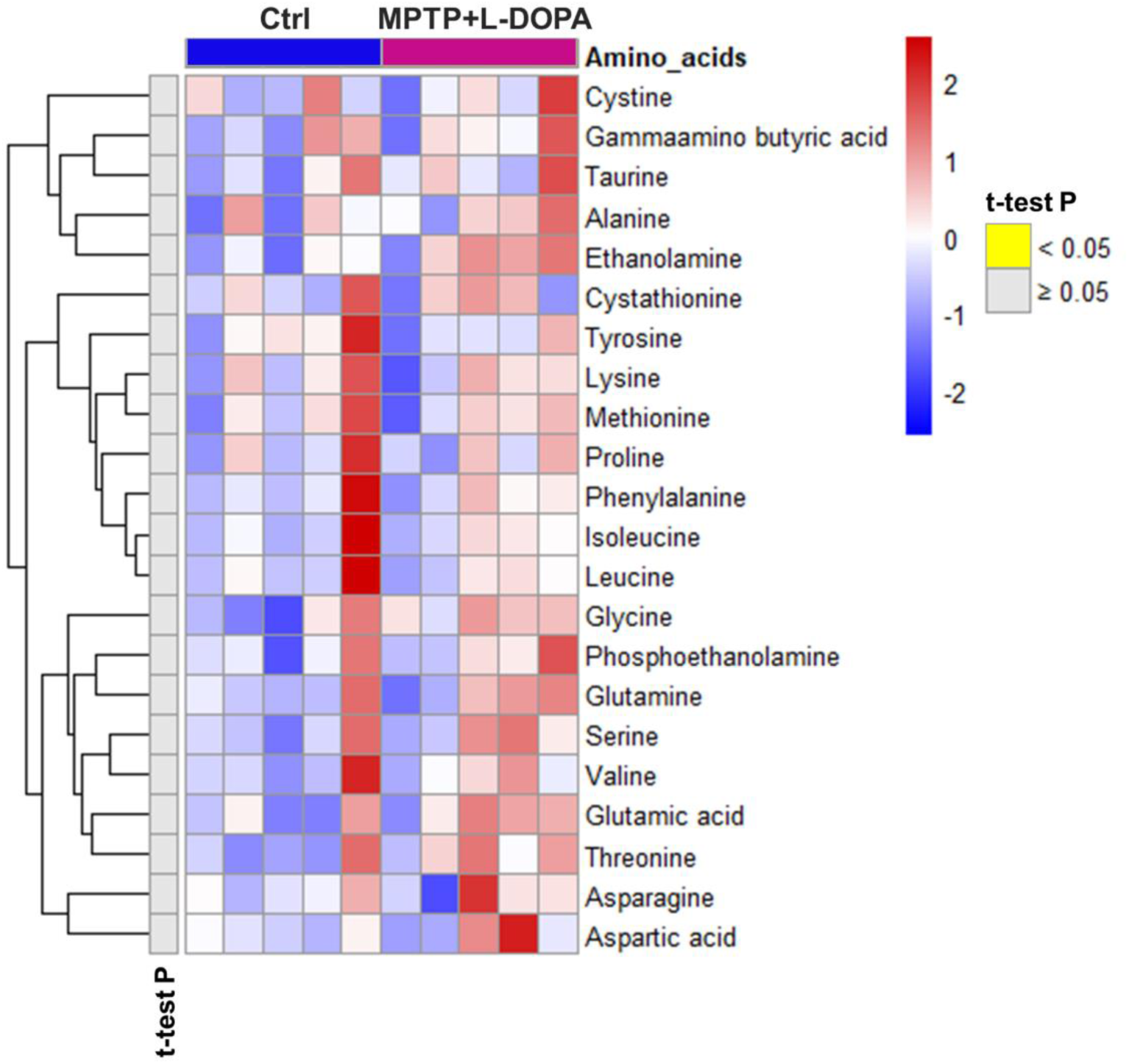
MPTP-intoxicated, L-DOPA-treated monkeys showed unaltered amino acid concentrations in the *post-mortem* superior frontal gyrus. The clustergram displays amino acids within the Monkeys’ SFG dataset. Rows represent amino acids (variables), while columns the samples, divided into Ctrl (n=5) and MPTP+L-DOPA (n=5). The elements of the clustergram illustrate amino acid concentrations in samples, standardized using a z-score, visualized using a color scale ranging from blue (lower) to red (higher). The first column indicates t-test statistical significance, showing not significant amino acids in gray. Abbreviations: Ctrl, control; MPTP, 1-methyl-4-phenyl-1,2,3,6-tetrahydropyridine; SFG, superior frontal gyrus.

### Upregulation of serine and proline and downregulation of phosphoethanolamine in the post-mortem caudate-putamen of Parkinson’s disease patients

We next investigated whether the pathological amino acid alterations observed in the putaminal samples of MPTP-intoxicated, L-DOPA-treated monkeys were also present in the *post-mortem* CPu from PD patients at different Braak LB stages (3–4 vs 6) compared to non-demented controls. To this end, we first performed univariate analyses in *post-mortem* Human CPu samples. A two-class comparison was initially conducted, assessing FC and t-test p-values between the PD and non-demented controls. This analysis revealed that three amino acids were significantly dysregulated in the PD group: serine (FC = 1.37; p-value 0.014) and proline (FC = 1.28; p-value 0.04) were upregulated, while phosphoethanolamine was downregulated (FC = –1.51; p-value 0.049).

Next, we performed a second analysis to include PD patients at different Braak LB stages (3–4 vs 6) and compared these groups to non-demented controls. To assess differential amino acid expression across the groups, a one-way ANOVA was performed, revealing a significant variation in serine levels (F2-29: 8.013; p-value: 0.0016) (Fig. 3A). A Tukey’s HSD post hoc test was conducted, revealing significant variation in serine levels (FDR 0.046), with an increase in Braak LB stage 6 compared to non-demented controls, and in Braak LB stage 3–4 compared to non-demented controls (Fig. 3B). Subsequent two-class comparisons between the Braak LB stage groups and non-demented controls highlighted further alterations in amino acid profiles of PD patients in the *post-mortem* CPu. A two-sample t-test was employed to assess statistical significance, while FC analysis was used to quantify the degree and direction of alterations in amino acid levels. The comparison between PD patients at Braak LB stage 3–4 and non-demented controls showed a significant upregulation of serine (FC 1.29; p-value 0.039) (Fig. 3C). When comparing the group of PD patients at Braak LB stage 6 to non-demented controls, we observed the upregulation of serine (FC 1.45; p-value 0.005), a concurrent increase in proline levels (FC 1.31; p-value 0.033), and a significant downregulation of phosphoethanolamine (FC –1.54; p-value 0.046) (Fig. 3D). Lastly, the comparison between the two groups of PD patients at different Braak LB stages (3–4 vs. 6) revealed a significant downregulation of arginine (FC –1.29; p-value 0.03) in the Braak LB stage 6 group relative to the stage 3–4 group (Fig. 3E).

**Figure 3.**
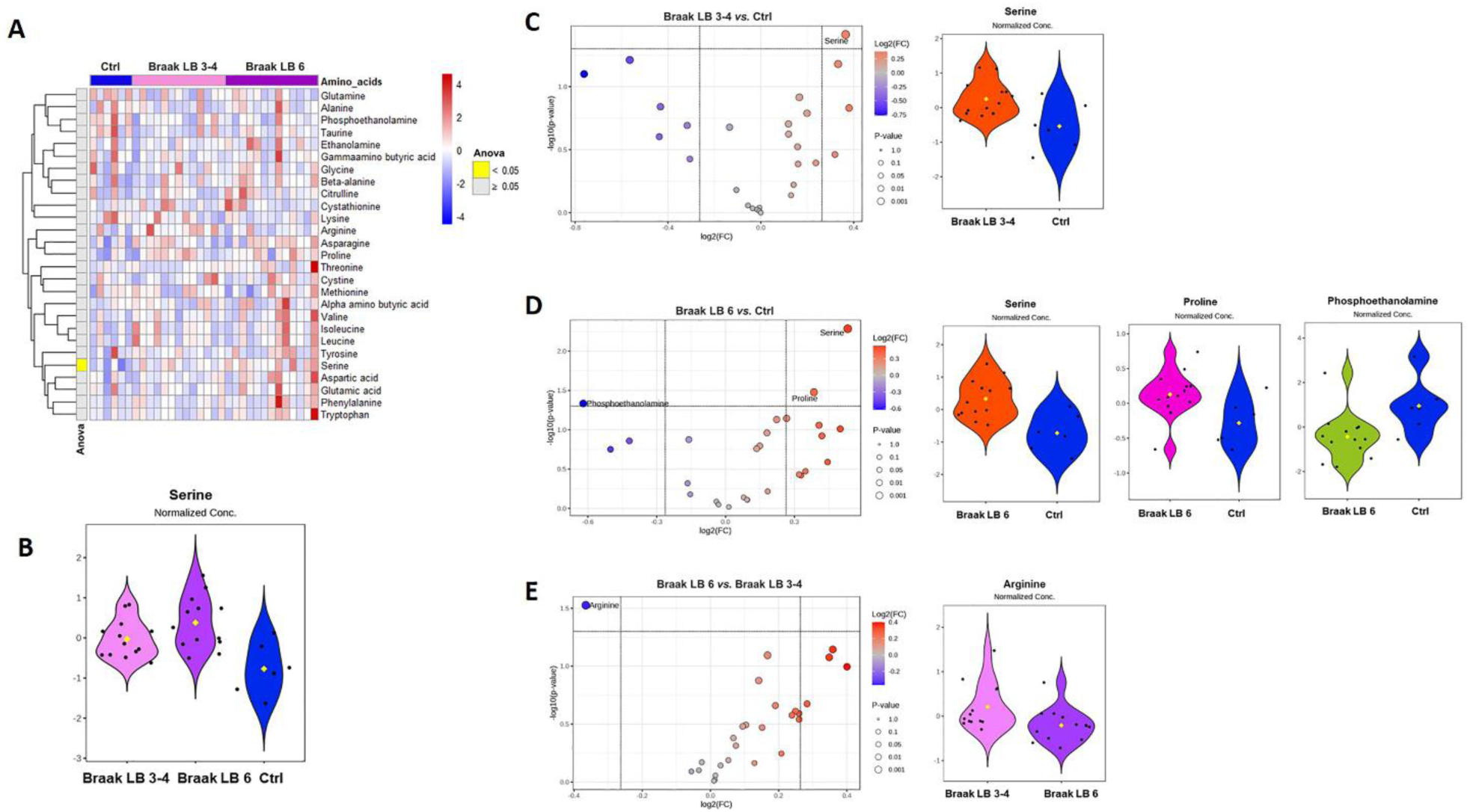
Serine, proline, and phosphoethanolamine levels are altered in the *post-mortem* caudate-putamen of Parkinson’s disease patients across Braak LB stages. A) The clustergram displays amino acids within the Human caudate-putamen dataset. Rows represent amino acids (variables), while columns the samples, divided into Ctrl (n=6), Braak LB 3-4 (n=13), Braak LB 6 (n=13). The elements of the clustergram illustrate amino acids concentrations in samples, standardized using a z-score, visualized using a color scale ranging from blue (lower) to red (higher). The first column indicates ANOVA statistical significance, showing the significant amino acids in yellow and not significant ones in gray. **B)** The violin plot displays the concentration of serine in Ctrl, and PD patients at Braak LB 3-4 and 6. The Tuckey’s HSD post hoc highlighted Braak LB 6 vs. Ctrl and Braak LB 3-4 vs. Ctrl as statistically significant comparisons (FDR 0.046). **C)** Left: The log2 (FC) and –10log (p-value) volcano plot illustrates the Braak LB 3-4/Ctrl ratio for each amino acid. Red points identify upregulated metabolites, while blue downregulated ones. Right:Violin plot of the significantly differential result is displayed. **D)** Left: The log2 (FC) and –10log (p-value) volcano plot illustrates the Braak LB 6/Ctrl ratio for each amino acid. Red points identify upregulated metabolites, while blue downregulated ones. Right: Violin plots of the significantly differential results are displayed. **E)** Left: The log2 (FC) and –10log (p-value) volcano plot illustrates the Braak LB 6/Braak LB 3-4 ratio for each amino acid. Red points identify upregulated metabolites, while blue downregulated ones. Right: Violin plot of the significantly differential result is displayed. Abbreviations: Ctrl, control; FC, fold change; LB, Lewy body.

### Amino acids levels are unaltered in the post-mortem superior frontal gyrus of PD patients

Next, we examined whether the amino acid alterations observed in the *post-mortem* CPu of PD patients were also present in cortical regions, such as the SFG, or whether they were region-specific, as previously observed in Parkinsonian monkeys. A univariate analysis was conducted across the *post-mortem* SFG samples of PD patients and non-demented controls (Human SFG dataset) with the aim of identifying differentially expressed amino acids in this specific brain region. The analysis was performed comparing the amino acid profiles of PD patients at Braak LB stages 4-5 and 6 against those of non-demented controls. A one-way ANOVA was conducted to evaluate differences in amino acid levels across the three groups, revealing no statistically significant alterations (Fig. 4A). This result was further confirmed by conducting a series of two-class comparisons, which also failed to identify any significant changes in amino acid levels between the groups (Fig. 4B-D). Importantly, the comparison of amino acid alterations found in striatal samples from PD patients and MPTP-intoxicated, L-DOPA-treated monkey results revealed a common significant upregulation of serine levels compared to their respective controls (Fig. 5).

**Figure 4.**
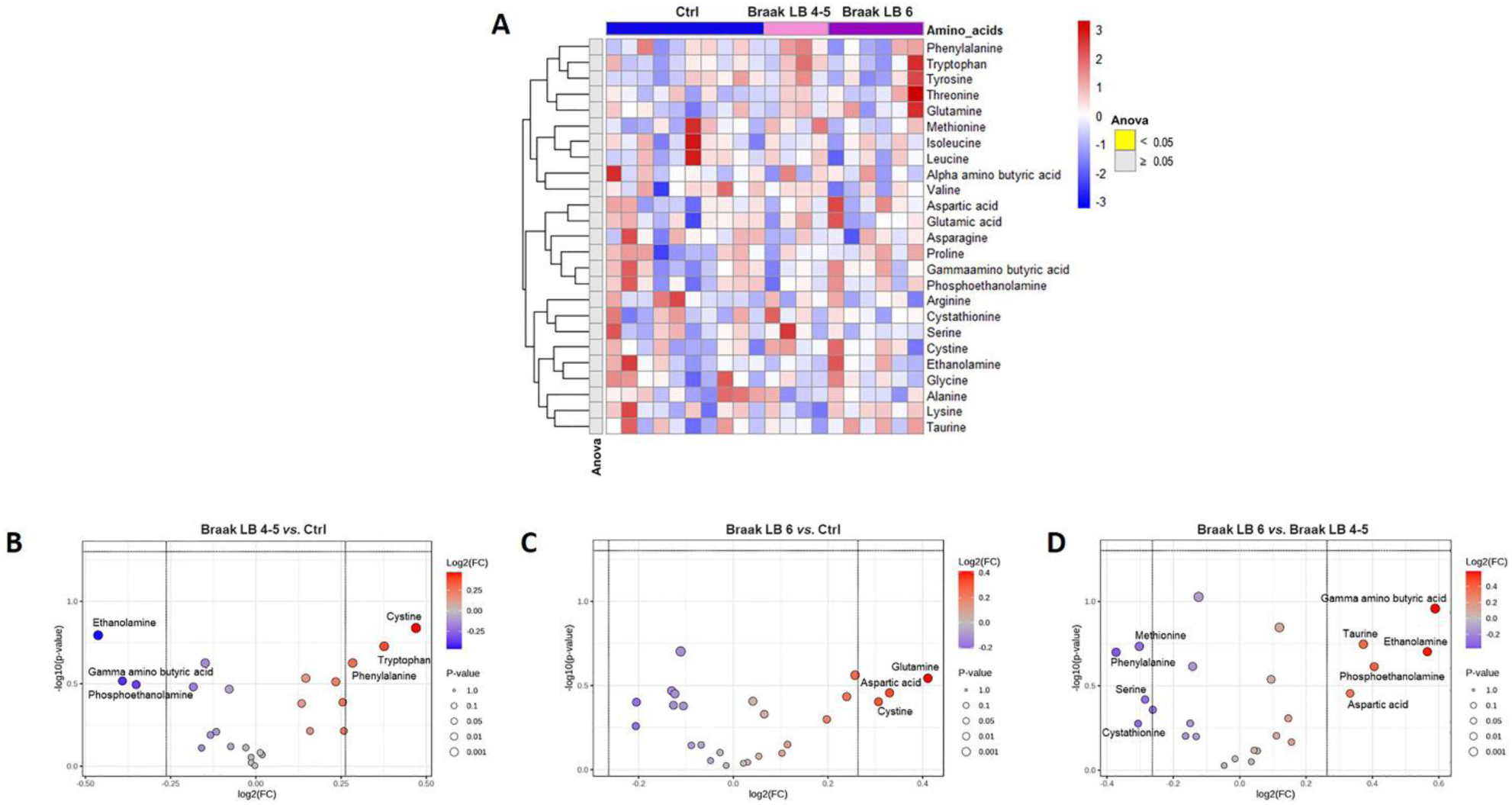
Parkinson’s disease did not induce variation amino acid levels in the superior frontal gyrus. **A)** The clustergram displays amino acids within the Human SFG dataset. Rows represent amino acids (variables), while columns the samples, divided into Ctrl (n=10), Braak LB 4-5 (n=4), Braak LB 6 (n=6). The elements of the clustergram illustrate amino acid concentrations in samples, standardized using a z-score, visualized using a color scale ranging from blue (lower) to red (higher). The first column indicates ANOVA statistical significance, showing not significant amino acids in gray. **B)** The log2 (FC) and –10log (p-value) volcano plot illustrates the Braak LB 4-5/Ctrl ratio for each amino acid. Red points identify upregulated metabolites, while blue downregulated ones, showing not significant amino acids. **C)** The log2 (FC) and –10log (p-value) volcano plot illustrates the Braak LB 6/Ctrl ratio for each amino acid. Red points identify upregulated metabolites, while blue downregulated ones, showing not significant amino acids. **D)** The log2 (FC) and –10log (p-value) volcano plot illustrates the Braak LB 6/Braak LB 4-5 ratio for each amino acid. Red points identify upregulated metabolites, while blue downregulated ones, showing not significant amino acids. Abbreviations: Ctrl, control; FC, fold change; LB, Lewy body; SFG, superior frontal gyrus.

**Figure 5.**
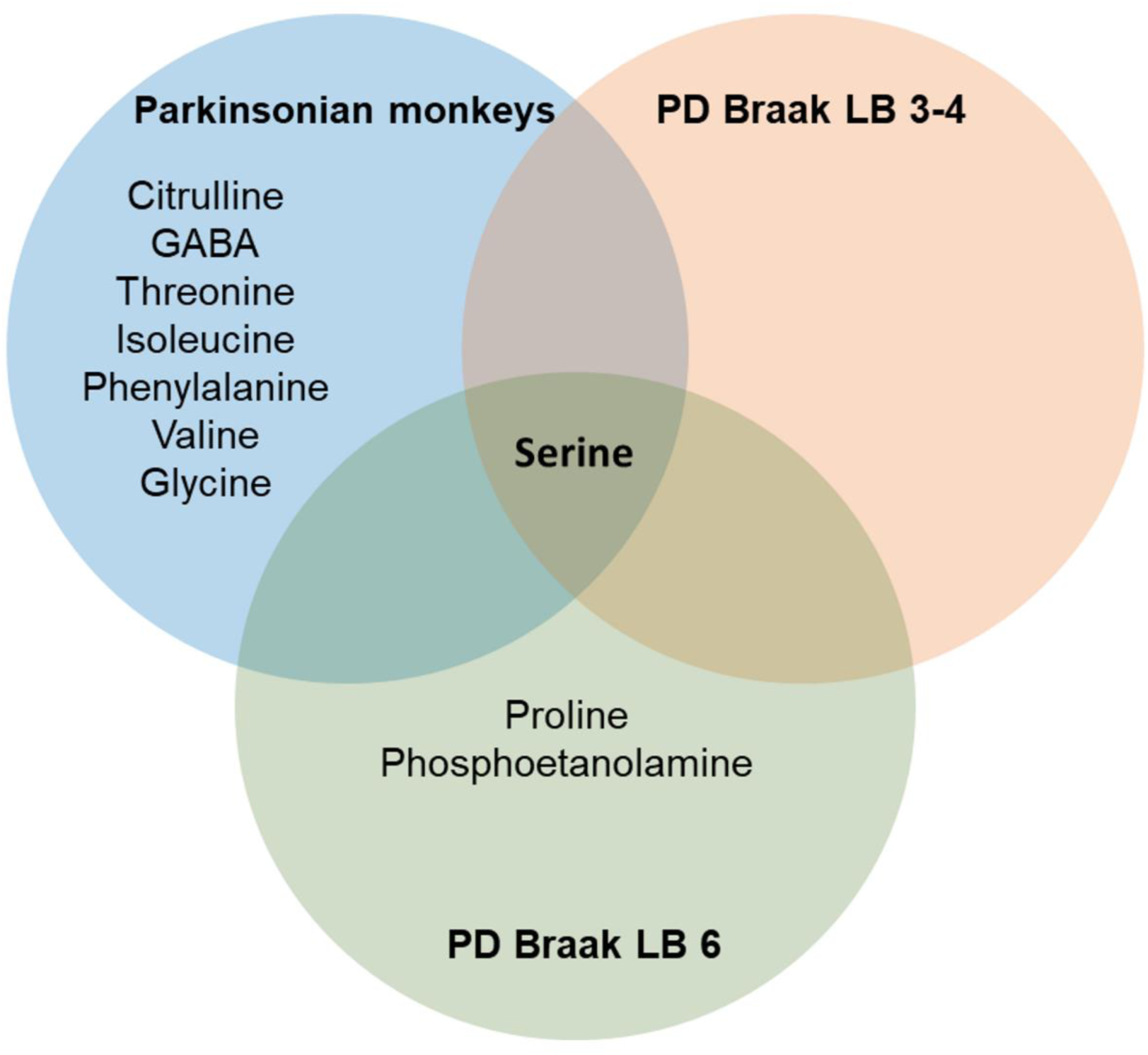
Common and distinct amino acid variations in the *post-mortem* striatal samples of PD patients and MPTP monkey models. Venn diagram of the results obtained in striatal *post-mortem* samples from Parkinsonian monkeys and PD patients at different Braak LB stages highlighted variation in serine levels as common feature, two exclusive altered amino acids for PD patients at Braak LB stage 6, while seven amino acids altered exclusively in Parkinsonian monkeys.

### Differential Amino Acid Expression in the Superior Frontal Gyrus of Alzheimer’s Disease Patients

Lastly, considering the role of the superior frontal gyrus in the pathophysiology of AD [40–42] and the absence of alterations observed in the PD cohort, we investigated whether cortical amino acid changes differ significantly across distinct neurodegenerative conditions. To address this, we performed a univariate analysis on the *post-mortem* SFG samples from AD patients and non-demented controls (Human SFG dataset). A t-test was conducted to assess statistical significance, and a FC analysis was performed to determine differences in amino acids levels. The results showed a net difference in amino acid concentrations across the two groups (Supplementary Fig. 1). The levels of five amino acids showed significant changes in AD patients compared to non-demented controls (Fig. 6A): Tryptophan (FC 1.37; p-value 0.002), phenylalanine (FC 1.34; p-value 0.003), threonine (FC 1.32; p-value 0.013), tyrosine (FC 1.27; p-value 0.022) and methionine (FC 1.32; p-value 0.024) were found to be upregulated in the AD group (Fig. 6B).

**Figure 6.**
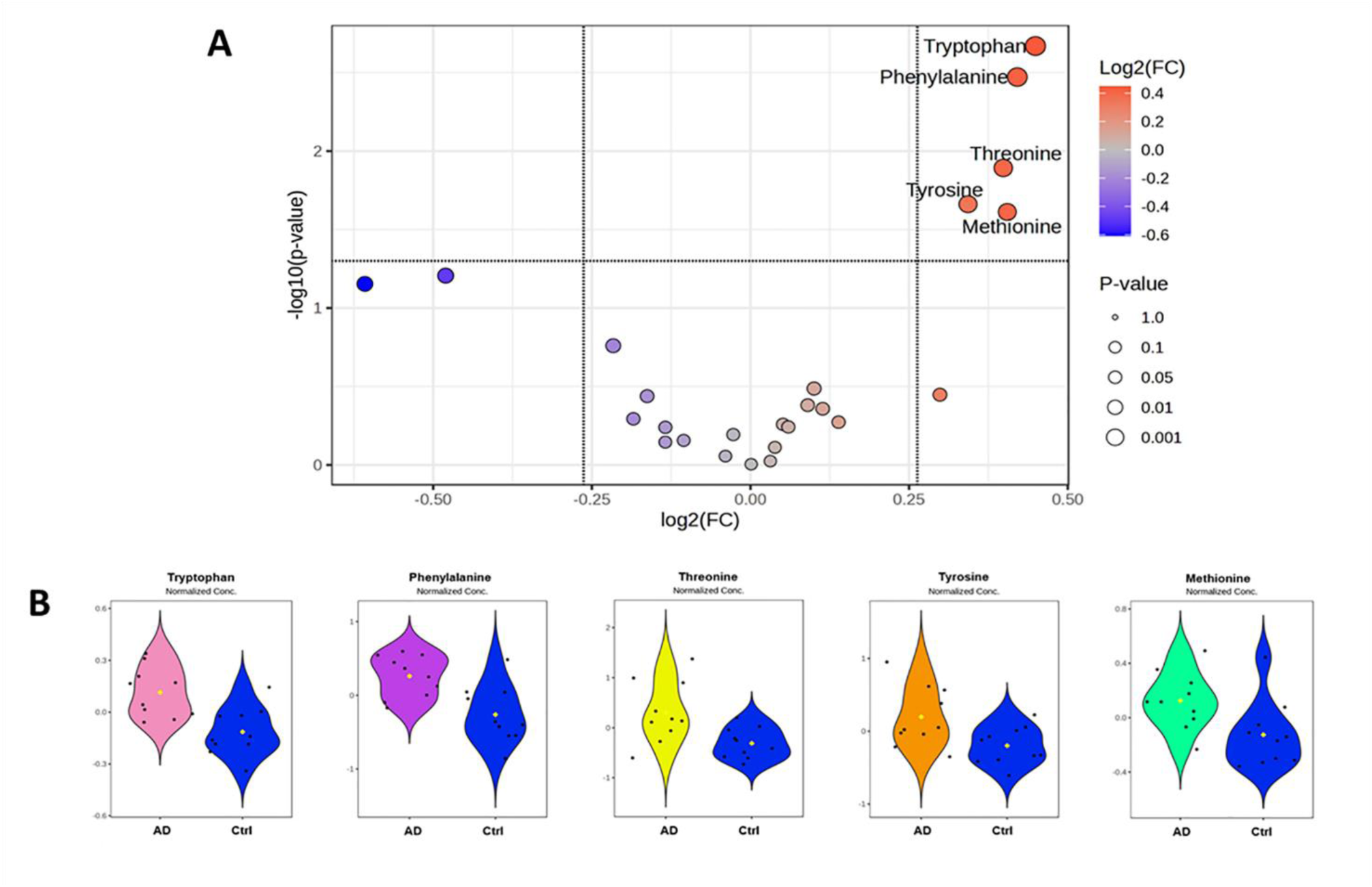
Alzheimer’s disease induces upregulation of tryptophan, phenylalanine, threonine, tyrosine and methionine in the *post-mortem* superior frontal gyrus. **A)** The log2 (FC) and –10log (p-value) volcano plot illustrates the AD/Ctrl ratio for each amino acid. Red points identify upregulated metabolites, while blue downregulated ones. **B)** Violin plots of the significantly differential result are displayed. Abbreviations: AD, Alzheimer’s disease; Ctrl, control; FC, fold change; LB, Lewy body.

## 4. Discussion

In the present study, we employed targeted UPLC-MS—a state-of-the-art analytical method renowned for its elevated sensitivity, selectivity, and broad dynamic range—to quantify the concentrations of 44 distinct amino acids in *post-mortem* brain samples from MPTP-treated monkeys, patients with PD or AD, and their respective healthy controls. The primary objective was to determine whether the metabolic alterations previously reported in the blood of PD patients, by using metabolomics and HPLC determinations [7,14,22,23,29,43], are also detectable in the CPu, a basal ganglia nucleus implicated in PD, and to evaluate the impact of neurodegenerative pathology on these central metabolic changes. Our results demonstrate distinct amino acid alterations associated with PD, specifically in the putamen of L-DOPA-treated, MPTP-intoxicated monkeys and in the CPu of PD patients at different Braak LB stages (3–4 and 6), compared to their respective controls. Notably, these alterations appear more circumscribed than those previously identified in the PD serum [7,14,23,29,43] and are region-specific, as they were not observed in another cortical brain region, the SFG. Importantly, comparative analyses between Parkinsonian monkeys and PD patients revealed that serine is the only amino acid consistently altered in both monkeys and humans. Lastly, by analyzing the SFG of patients with AD and their respective controls, we provide evidence highlighting the impact of neurodegenerative pathology on cortical amino acid level alterations.

Accumulating studies have revealed significant metabolic dysregulation in the peripheral biofluids— particularly plasma and serum—of PD patients relative to healthy individuals [13–15,20–24,43]. These alterations encompass dysregulation in amino acid metabolism, lipid homeostasis, mitochondrial energy metabolism, and purine biosynthesis pathways [7,14,29], highlighting the multi-organ nature of PD [44–47].

While these studies provide key insights into the blood metabolic state of the disease, it remains uncertain whether these alterations reflect modifications in central biochemical processes— particularly within the basal ganglia—or are systemic in nature. As highlighted by a recent meta-analysis revealing inconsistencies across 74 metabolic studies in PD patients [29], clarifying the origin of these alterations is crucial for validating diagnostic and prognostic biomarkers in PD.

To address this issue, the present study employs targeted UPLC-MS to precisely quantify in the *post-mortem* brain samples of MPTP-intoxicated monkeys, patients with PD or AD, and relative controls, the central levels of the 44 amino acids previously analysed in the sera of PD patients and controls using the same analytic approach [14].

In the *post-mortem* putamen of L-DOPA-treated, MPTP-intoxicated monkeys, UPLC-MS analysis revealed significant perturbations in the levels of eight metabolites compared to control monkeys. This perturbation included: GABA, citrulline, threonine, isoleucine, phenylalanine, valine, glycine, and serine, all of which exhibited upregulated concentrations.

In line with our findings, previous studies have shown higher GABA levels in the striatum of MPTP-treated monkeys [48], as well as within the basal ganglia of patients with PD [49]. Considering that the striatum is made up for about the 90% of GABAergic medium spiny neurons [50], whose activity is regulated by nigrostriatal dopamine inputs [51–55], we argue that the upregulation of GABA biosynthesis might represent a compensatory mechanism in response to dopamine depletion and L-DOPA treatment, potentially aiming to restore the delicate balance of neuronal activity.

Previous reports have indicated reduced levels of citrulline in the sera [43] and CSF [56] of PD patients compared to controls. In contrast, to the best of our knowledge, alterations in the levels of citrulline in the basal ganglia of PD patients remain unexplored. The observed increase in citrulline, an intermediate in the urea cycle and a precursor of arginine and nitric oxide [57], found in the putamen of Parkinsonian monkeys suggest alterations in mitochondrial function and nitrosative stress, both of which are well-established features of PD pathology [58,59]. Furthermore, upregulation of citrulline may impact the post-translational citrullination/deamination process, as recently evidence in a prodromal rodent model of PD [60].

Similarly, the elevated levels of branched-chain amino acids (BCAA) isoleucine and valine, the aromatic amino acid phenylalanine, and the essential amino acid threonine, could reflect altered protein metabolism, mitochondrial dysfunction, and neurotransmitter synthesis. While the role of BCAA in PD pathogenesis is still debated [61–63], a large-scale study using UK Biobank data found that higher blood BCAA levels were associated with reduced risk of dementia and AD, but potentially increased risk of PD [64]. Remarkably, in our recent serum metabolomics study comparing PD patients to healthy controls, threonine and valine emerged as key discriminatory metabolites [14].

Furthermore, the upregulation of glycine and serine is of considerable interest given their roles in neurotransmission and their involvement in various metabolic pathways, including antioxidant defence [65–67]. In this regard, findings from preclinical animal models and PD patients suggest that stimulation of the GluN1 subunit of glutamatergic N-methyl-D-aspartate (NMDA) receptors—where both glycine and D-serine bind—may confer therapeutic benefits in PD [68–71].

In contrast to the putamen, UPLC-MS analysis in SFG—a cortical region receiving dopaminergic innervation from the ventral tegmental area but not the substantia nigra pars compacta [72]—showed no significant changes in amino acid levels between L-DOPA-treated, MPTP-intoxicated monkeys and control monkeys. These findings suggest that the amino acid metabolic alterations observed in PD brains are region-specific, predominantly affecting areas most impacted by dopaminergic denervation, such as the putamen.

The pronounced alterations in amino acid levels observed in the *post-mortem* putamen of Parkinsonian monkeys prompted us to investigate whether similar changes were present in the same brain region of PD patients, stratified by Braak Lewy body stages (3–4 and 6). In contrast to our observations in Parkinsonian monkeys, UPLC-MS analysis in the *post-mortem* CPu of the entire cohort of PD patients revealed a notable alteration in only three amino acids— serine, phosphoethanolamine, and proline —compared to non-demented controls. Interestingly, increased levels of phosphoethanolamine and proline have been previously reported in the sera of PD patients [43,73], while the upregulation of serine is consistent with our earlier HPLC and metabolomic study conducted on *post-mortem* CPu and serum samples from PD patients [14,21].

The difference between patients with PD and Parkinsonian monkeys in amino acid level variations can be attributed to several factors. These include variations in the duration of treatment with anti-Parkinsonian medications (4-5 months in monkeys versus years in PD patients), post-mortem interval (seconds-to-minutes in monkeys vs. hours in humans) sex-related heterogeneity, differential peripheral organ dysfunctions [74,75] and the divergent aetiologies leading to nigrostriatal dopaminergic neuron degeneration (toxin-induced in monkeys versus multifactorial in idiopathic PD humans).

Interestingly, upon patient stratification by Braak LB stages, serine was the only amino acid found upregulated in patients at stages 3–4, whereas at stage 6, serine and proline were significantly upregulated while phosphoethanolamine was downregulated relative to non-demented controls.

When comparing PD patients at Braak LB stages 3–4 with those at stage 6, we found that arginine levels were significantly downregulated in patients at stage 6. Arginine is critically involved in the synthesis of nitric oxide, an important regulator of physiological processes in the brain including synaptic plasticity, intracellular signalling, and cerebral blood flow [76]. Although the functional consequences of arginine reduction within the *post-mortem* CPu PD samples remain to be fully explored, it is noteworthy that repeated systemic pre-treatment with L-arginine has been shown to prevent MPTP-induced dopaminergic cell loss in the substantia nigra pars compacta in mice [77].

When we extended the UPLC-MS analysis to the *post-mortem* SFG, we failed to observe any statistically significant differences in the concentration of the 44 amino acids analyzed between patients with PD and non-demented controls, a finding consistent with our results in Parkinsonian monkeys.

The present metabolomic analysis on the brain of PD patients highlight a key aspect when compared with prior meta-analysis in the serum/plasma of patients with PD [7,23,29,43]. Whereas these studies have identified several amino acids alterations that distinctly separated PD patients from controls, alterations in the *post-mortem* CPu were restricted to only three amino acids. While we acknowledge that differences in sample size and subject characteristics between the present study and previous may have contributed to this discrepancy, we hypothesize that the extensive alterations detected in serum likely reflect multisystemic pathophysiological changes associated with the disease. In contrast, the limited amino acid alterations observed in the *post-mortem* CPu of PD patients may more specifically reflect central neuropathological features of the disease. Nonetheless, further investigation involving *post-mortem* CPu samples from larger cohorts of male and female PD patients, alongside the integration of other OMIC techniques, is required to clarify this relevant issue and to gain further insights into the biological relevance of central and peripheral metabolic biomarkers.

Notably, across all amino acids found to be altered in the putamen of Parkinsonian monkeys, as well as in the serum [14,22] and CPu of PD patients, serine is the only metabolite that consistently emerges. In this regard, it is noteworthy that serine exists in two distinct enantiomeric configurations, D– and L-serine, whose brain concentrations are strictly regulated and balanced due to their critical central effects. Specifically, while in the brain the atypical amino acid D-serine acts as a co-agonist of the NMDA receptor [67,78], L-serine serves as a metabolic precursor for sphingolipids, purine nucleotides, antioxidants, and NMDA receptor co-agonists (glycine and D-Serine) [79,80]. Although UPLC-MS analysis does not allow for the discrimination of serine enantiomers, our previous HPLC studies demonstrated elevated levels of both D and L-serine in brain tissue from experimental PD models, *post-mortem* brain samples from PD patients, and CSF from *de novo* PD patients compared to controls [21,81]. These findings suggest that the increase in serine observed in the *post-mortem* CPu of PD patients in the present *post-mortem* study likely reflects a concurrent elevation of both enantiomers. In particular, we hypothesize that elevation in CPu D-serine levels may reflect an adaptative NMDAR-related response aim at contrasting the progressive loss of striatal dopamine signalling in PD. This hypothesis is supported by recent evidence demonstrating that D-serine modulates dopamine release in the CPu by enhancing glutamatergic transmission through NMDA receptors on dopaminergic nigrostriatal neurons [82]. Moreover, elevation of spinal D-serine levels, achieved through either genetic manipulation or exogenous D-serine administration, was found to attenuate motor symptoms, improve blood–brain barrier (BBB) function and permeability, and counteract neuroinflammation in the experimental autoimmune encephalomyelitis (EAE) mouse model of multiple sclerosis [83]. These findings are highly relevant, given that BBB disruption and neuroinflammation are well-established neuropathological mechanisms contributing to PD [84,85].

While further investigations are needed to clarify this clinical issue, current data suggest that target amino acid supplementation may represent a possible therapeutic option for symptom relief by addressing underlying biochemical imbalances.

Lastly, we evaluated by UPLC-MS the amino acid metabolic profile in *post-mortem* SFG samples from controls and AD patients—a brain region known to be affected by this neurodegenerative disorder [40–42]. Through this approach, we aimed to evaluate the impact of different neurodegenerative conditions on cortical amino acid metabolism. Consistent with the known relevance of this brain region in AD, and in stark contrast to our findings in PD patients, we observed significant alterations in amino acid metabolism in the *post-mortem* SFG of patients with AD. Specifically, our data indicated a net upregulation in the levels of tryptophan, phenylalanine, threonine, tyrosine, and methionine in AD patients compared to controls. These results align with previous reports documenting alterations in the metabolism of these amino acids in brain regions highly susceptible to AD pathology, such as the hippocampus [86], the Brodmann area 9—which is located within the SFG [86,87]—and the temporal cortex [88]. Notably, these findings underscore that amino acid level alterations in cortical regions such as the SFG are highly dependent on the underlying neuropathological condition, as clearly demonstrated by the divergent metabolic profiles observed in patients with PD and AD compared to controls.

The primary limitation of this study is the imbalance between male and female subjects—both in animal models and human cohorts—and the inability to stratify patients by genetic background, likely restricting the cohort to idiopathic cases due to the limited sample size. Such constraints arise from the inherent difficulty of obtaining *post-mortem* brain tissue samples from the study population. The study’s primary strength lies in stratifying the PD cohort by Braak LB stages (3–4 vs. 6), thus providing direct insight into disease progression, and in employing targeted UPLC-MS for amino acid profiling—a platform frequently used in clinic renowned for its high sensitivity, selectivity, and quantitative accuracy in complex biological matrices.

In summary, the present UPLC-MS study, which profiled 44 amino acids in *post-mortem* striatal and cortical samples from L-DOPA-treated, MPTP-intoxicated monkeys and PD patients at various disease stages, revealed that metabolic alterations were numerically less widespread compared to those previously reported in the PD serum [7,23,29,43] and confined to the CPu, with serine being uniquely upregulated across both species and disease stages. In contrast, the SFG remained metabolically unaltered in PD but exhibited distinct amino acid dysregulation in AD, thus demonstrating both disease– and region-specific metabolic signatures. In conclusion our findings underscore a potential central role for serine accumulation in nigrostriatal pathology and highlight its potential as a biomarker for PD. Furthermore, our results emphasize the critical importance of differentiating between central and peripheral metabolic changes to advance our understanding of PD pathophysiology, improve the validity of biochemical biomarkers, and facilitate the development of novel therapeutic strategies for this neurodegenerative disorder.

## Acknowledgements

Post-mortem human caudate putamen and superior frontal gyrus samples were provided by The Netherlands Brain Bank (Netherlands Institute for Neuroscience, Amsterdam, open access: www.brainbank.nl). We acknowledge Alessia Casamassa, Mattia Miroballo, Giorgia Donati, Giulia Sansone, Giada Torresi and Martina Garofalo for their technical support.

## Author contributions

**Jacopo Gervasoni**: Data curation, Formal analysis, Investigation **Anna Di Maio**: Investigation **Marcello Serra** Writing – original draft, Writing – Critical revision & editing. **Lavinia Santucci**: Investigation **Michela Cicchinelli:** Data curation, Formal analysis **Tommaso Nuzzo:** Critical revision & editing. **Qin Li**: Investigation, **Marie-Laure Thiolat**: Investigation, **Micaela Morelli:** Critical revision. **Francesco Errico**: Critical revision & editing. **Andrea Urbani:** Data curation, Formal analysis, **Erwan Bezard**: Critical revision & editing. **Alessandro Usiello:** Conceptualization, Data curation, Funding acquisition, Project administration, Supervision, Visualization, Writing – original draft, Writing – Critical review and editing.

## Availability of data and materials

The raw data supporting the findings of the present study are available on request from the corresponding author

## Declaration

### Ethics approval and consent to participate

The post-mortem samples included in the present study were purchased from the Netherlands Brain Bank. All material and data acquired by the Netherlands Brain Bank adhere to a strict protocol requiring written informed consent. In instances where individuals are unable to provide informed consent due to health conditions, authorization is obtained from their legal representative, in accordance with the Netherlands Civil Code (Burgerlijk Wetboek). Additional details are available at: https://www.brainbank.nl/about-us/ethics/

## Consent for publication

Not applicable.

## List of abbreviations

α-syn: Alpha-synuclein
AD: Alzheimer’s disease
ANOVA: Analysis of Variance
BCAA: Branched-chain amino acids
BBB: Blood–brain barrier
CPu: Caudate-putamen
CSF: Cerebrospinal fluid
EAE: Experimental autoimmune encephalomyelitis
FC: Fold change
FDR: False Discovery Rate
GABA: Gamma-aminobutyric acid
HPLC: High-Performance Liquid Chromatography
HSD: Honestly Significant Difference
LB: Lewy body
m.v.: Missing values
MPTP: 1-methyl-4-phenyl-1,2,3,6-tetrahydropyridine
NMDA: N-methyl-D-aspartate
PD: Parkinson’s disease
PMI: Post-mortem interval
PPCA: Probabilistic Principal Component Analysis
SFG: Superior frontal gyrus
TCA: Trichloroacetic acid
UPLC-MS: Ultra-Performance Liquid Chromatography-Mass Spectrometry

