## Supplementary Figure 1 for "Ultra-performance liquid chromatography-mass spectrometry analysis of post-mortem brain tissue reveals specific amino acid profile dysregulation in Parkinson’s Disease and Alzheimer’s Disease patients"

**Supplementary Materials**

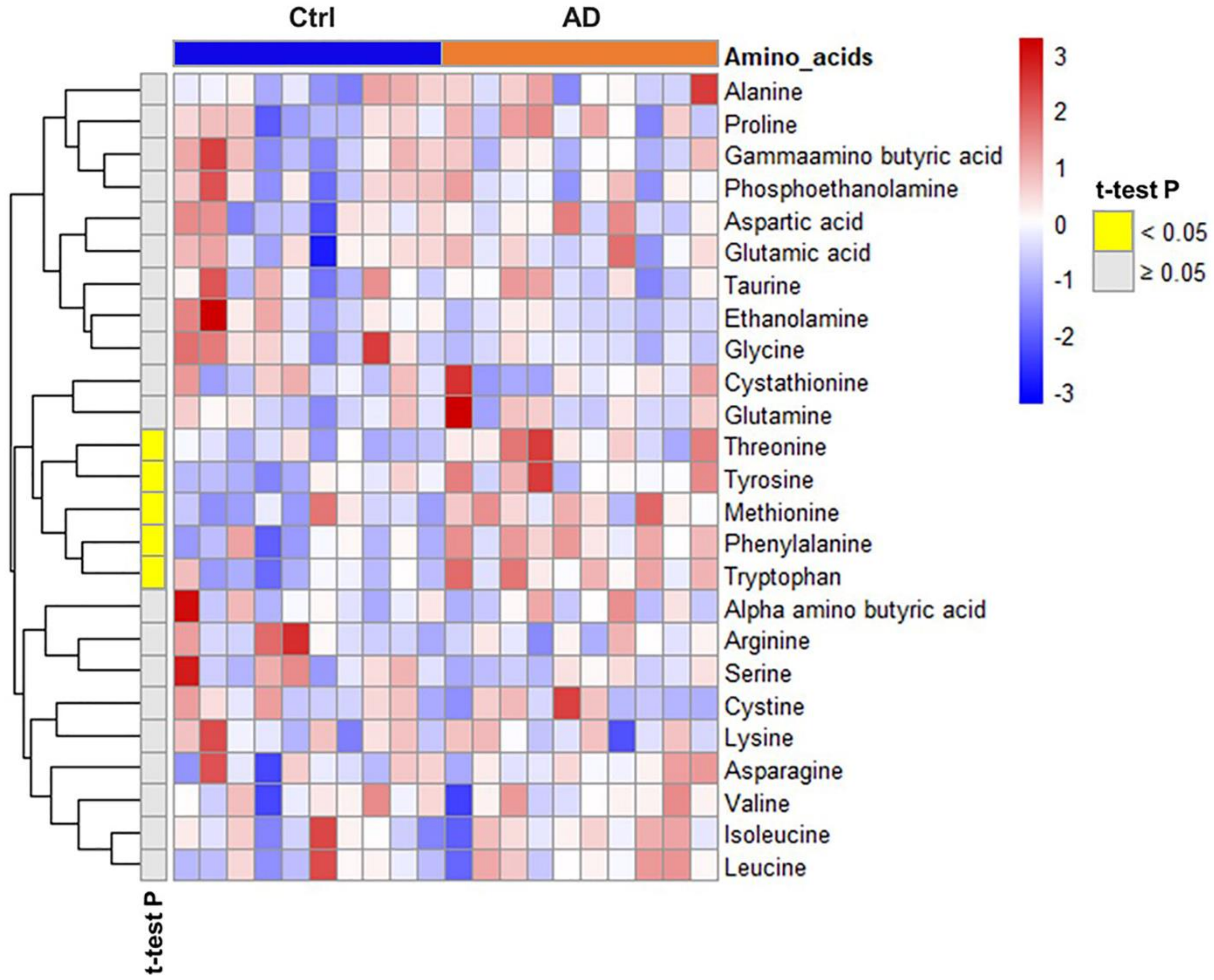


**Supplementary Figure 1. Alzheimer's disease induces upregulation of tryptophan, phenylalanine, threonine, tyrosine and methionine in the post-mortem superior frontal gyrus**. The clustergram displays amino acids within the Human SFG dataset. Rows represent amino acids (variables), while columns the samples, divided into AD (n=10) and Ctrl (n=10). The elements of the clustergram illustrate amino acid concentrations in the samples, standardized using a z-score, visualized using a color scale ranging from blue (lower) to red (higher). The first column indicates t-test statistical significance, showing the significant amino acids in yellow and not significant in gray.
